# Aligning sequences to general graphs in *O*(*V* + *mE*) time

**DOI:** 10.1101/216127

**Authors:** Mikko Rautiainen, Tobias Marschall

## Abstract

Graphs are commonly used to represent sets of sequences. Either edges or nodes can be labeled by sequences, so that each path in the graph spells a concatenated sequence. Examples include graphs to represent genome assemblies, such as string graphs and de Bruijn graphs, and graphs to represent a pan-genome and hence the genetic variation present in a population. Being able to align sequencing reads to such graphs is a key step for many analyses and its applications include genome assembly, read error correction, and variant calling with respect to a variation graph. Given the wide range of applications of this basic problem, it is surprising that algorithms with optimal runtime are, to the best of our knowledge, yet unknown. In particular, aligning sequences to cyclic graphs currently represents a challenge both in theory and practice. Here, we introduce an algorithm to compute the minimum edit distance of a sequence of length *m* to any path in a node-labeled directed graph (*V, E*) in *O*(*|V |*+*m*|*E*|) time and *O*(*|V |*) space. The corresponding alignment can be obtained in the same runtime using 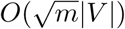 space. The time complexity depends only on the length of the sequence and the size of the graph. In particular, it does not depend on the cyclicity of the graph, or any other topological features.

## 1 Introduction

Aligning two sequences is a classic problem in bioinformatics. The standard dynamic programming (DP) algorithm, introduced by Needleman and Wunsch in 1970 [18], aligns two sequences of length *n* in *O*(*n*^2^) time. In 2015, it was shown that this time complexity is optimal in the sense that an *O*(*n*^2*−ϵ*^) algorithm for computing edit distance does not exist unless the strong exponential time hypothesis is false [2]. Countless variants of this classic DP algorithm exist, in particular its generalization to local alignment [24], where the alignment can be between any substrings of the two sequences, and semi-global alignment [23] where one sequence (*query*) is entirely aligned to a substring of the other (*reference*).

In addition to sequences, graphs whose nodes or edges are labeled by characters are commonly used for many applications in bioinformatics, for instance for genome assembly [13, 5] and multiple sequence alignment [10]. Currently, we witness a strong interest in the use of graphs also as a potential alternative to “linear” reference genomes [25, 19]. Graph-based reference genomes hold the promise of removing any reference bias and can naturally encode also complex variation. With an increasing usage of graphs, algorithms for aligning reads to graphs are also of growing interest and have already been applied successfully for purposes such as genome assembly [1] and error correction [21].

In this paper, we study the problem of semi-global sequence-to-graph alignment. The idea is that instead of a query sequence and a reference sequence (like in classic semi-global alignment), we have a query sequence and a directed, node-labeled graph as a reference. We then seek to find a path in the graph that has minimum edit distance to the query sequence.

### Related Work

*Partial order alignment* [11] (POA) is an extension of the standard DP for sequence alignment, where the input is a directed acyclic graph (DAG) and a sequence to be aligned to that graph. Each node in the DAG gives rise to a column in the DP matrix. Then, the DP recurrence takes into account each in-neighbor of a node and processes nodes in topological order. Assuming *V ∈ O*(*E*), this leads to a runtime of *O*(*mE*) where *m* is the length of the sequence and *E* is the number of edges.

Different approaches employing the seed-and-extend paradigm [4] to align short reads to DAGs have been proposed [22, 9] with the main goal of fast (yet heuristic) alignment suitable for applications. For aligning sequences to cyclic graphs, the graph can be “unrolled” into a DAG. Unrolling constructs a DAG whose paths of a given length *k* or smaller are all present in the cyclic graph, and vice versa. The result of this process is called a *k*-DAG [28]. The sequence is then aligned to the DAG using POA, and the result is mapped back into the cyclic graph. This is the approach used by the variation graph tool vg (https://github.com/vgteam/vg). For long reads, the *k* parameter has to be at least the length of the read. However, this method has the disadvantage that the graph size can grow drastically and finding the smallest *k*-DAG is NP-hard [28]. Unrolling is therefore unsuitable for aligning long reads to complex cyclic graphs.

V-align [28] is an alignment algorithm for cyclic graphs (and the winner of the best poster award at RECOMB 2017). It can be considered a generalization of the POA algorithm with a special method for handling cyclic nodes. It uses a linear ordering of the nodes and considers “in-order” and “out-of-order” nodes separately. The *in-order* nodes are nodes whose in-neighbors are all earlier in the ordering, whereas the “out-of-order” nodes have at least one in-neighbor later in the ordering. The runtime of V-align is *O*((*V′* + 1)*mE*) where *m* is the length of the sequence, *E* is the number of edges and *V′* is the number of out-of-order nodes in the used ordering. Note that *V′* is at least as large as the graph’s minimum feedback vertex set.

Limasset et al. propose a procedure for mapping reads to de Bruijn graphs [12], but it is heuristic in the sense that it is not guaranteed to find the optimal alignment. The genome assembler hybridSPAdes [1] aligns long reads to assembly graphs and shows that sequence-to-graph alignment can be re-phrased as a shortest path problem. Using Dijkstra’s algorithm leads to an algorithm that takes *O(|E|m* + *|V|m* log(*|V|m*)) time.

### Contributions

We present an algorithm for finding an optimal alignment between a sequence of length *m* and a directed node-labeled graph (*V, E*) in *O*(*V* + *mE*) time. Our algorithm works on any graph, including cyclic graphs, and the time complexity does not depend on the cyclicity or the shape of the graph. Since algorithms to compute edit distances in *O*(*n*^2*−ϵ*^) time are unlikely to exist [2], it is likewise unlikely that asymptotically faster algorithms to align a sequence of length *n* to a graph with *|V|,|E| ∈ O*(*n*) exist, because sequence-to-sequence alignment is a special case of sequence-to-graph alignment. We furthermore generalize observations on the maximum difference between cells in the DP table [27] to sequence-to-graph alignment. These insights might have utility for designing efficient implementations, for instance through bit-parallelism. Moreover, we show how to generalize sequence-to-graph alignment to affine gap costs, albeit at a slower runtime of *O*(*|E|m* + *|V |m* log *|V |*).

## 2 Semi-Global Sequence-to-Graph Alignment with Unit Costs

We make some definitions to establish notation and to formally state the semi-global sequence-to-graph alignment (SG^2^A) problems studied in this paper. For ease of exposition, we focus on the unit cost case first and generalize it affine gap costs in Section 3.

### Definition 1

(Sequence graph). We define a *sequence graph* as a tuple *G* = (*V, E, σ*), where *V* = *{v*_1_, …, *v*_*n*_} is a finite set of nodes, *E ⊂ V × V* is a set of directed edges and *σ*: *V →* Σ assigns one character from the alphabet Σ to each node. We refer to the sets of indices of in-neighbors and out-neighbors of node *v*_*i*_ as 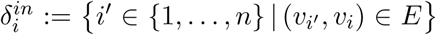 and 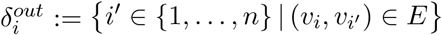, respectively.

### Definition 2

(Path sequence). Let *p* = (*p*_1_, …, *p*_*k*_) be a path in the sequence graph *G* = (*V, E, σ*); that is, *p_i_ ∈ V* for *i ∈* {1, …, *k*} and (*p*_*i*_, *p*_*i*+1_) *∈ E* for *i ∈* {1, …, *k* − 1}. Then, the *path sequence* of *p*, written *σ*(*p*), is given by *σ*(*p*_1_)*σ*(*p*_2_) … *σ*(*p*_*k*_).

We note that this definition of paths and path sequences includes the possibility of repeated vertices. With this definition, we can now state our main problem (unit cost SG^2^A) formally.

### Problem 1

(Unit Cost Semi-Global Sequence-to-Graph Alignment). Let a string *s ∈* Σ^*∗*^ and a sequence graph *G* = (*V, E, σ*) be given. Find a path *p* = (*p*_1_, …, *p*_*k*_) in *G* such that the edit distance *d*(*σ*(*p*), *s*) is minimized and report a corresponding alignment of *σ*(*p*) and *s*.

In the remainder of this paper, we assume an arbitrary but fixed string *s ∈* Σ^*∗*^ with *|s|* = *m* and sequence graph *G* = (*V, E, σ*) to be given. For now, we assume that *|V | ∈ O*(*|E|*); in a later section, we will justify this assumption. We approach Problem 1 by generalizing the standard dynamic programming (DP) algorithm for edit distance calculation. In our case, the DP matrix has one column per node *v_i_ ∈ V* and one row per character *s*_*j*_ from *s ∈* Σ^*∗*^. We seek to compute values *C*_*i,j*_ for *i ∈* {1, …, |*V* |} and *j ∈* {1, …, |*s*|} such that *C*_*i,j*_ is the minimum edit distance *d*(*p, s*[1..*j*]) over all paths *p* ending in node *v*_*i*_.

### Definition 3

(Recurrence for unit cost SG^2^A). Set *C*_*i*,1_ = ∆_*i,*1_ for all *i ∈* {1, …, |*V* |}. And define

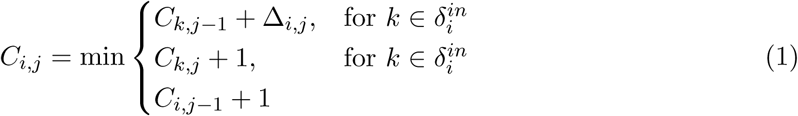

where ∆_*i,j*_ the mismatch penalty between node character *σ*(*v*_*i*_) and sequence character *s*_*j*_, which is 0 for a match and 1 for a mismatch.

The standard DP formulation for semi-global alignment is a special case of this recurrence, where the input graph is just a linear chain of nodes. Another special case of our formulation is *partial order alignment* [11] (POA), which aligns a sequence to a DAG and hence does not allow cyclic dependencies. For general graphs, which can contain cycles, we note that Recurrence (1) can likewise contain cyclic dependencies, where the value *C*_*i,j*_ depends on value *C*_*x,j*_ which in turn depends on *C*_*i,j*_. Therefore, it is not immediately obvious that it is well-defined and leads to the intended semantics of the DP cells. Establishing this is the goal of the next section.

### 2.1 Existence and Uniqueness

Recurrence 1 cannot be directly used to calculate the scores for cyclic regions. Instead, we consider this recurrence a constraint on the cell scores, and then find an assignment of scores to the cells which satisfies the recurrence. There exists exactly one assignment of scores that satisfies the recurrence, which we prove in the following. To this end, we employ a classic relationship between edit distances and shortest path problems: we define an *alignment graph* with one node for each cell in our DP table, such that the distance from a virtual source node equals the value in the corresponding cell [14], as illustrated in Figure 1(right). Formally, the alignment graph (not to be confused with a sequence graph) is defined as follows.

**Figure 1:**
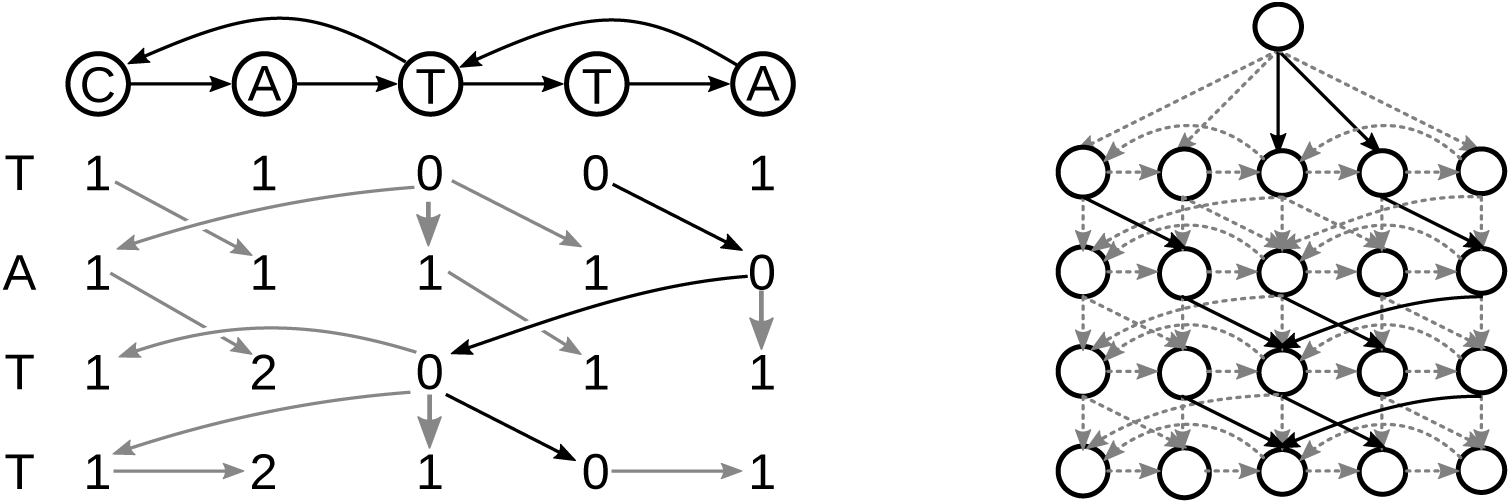
Left: the DP matrix of the alignment between a sequence graph (top) and a sequence (left). The gray arrows are the backtrace and the solid black arrows show the optimal alignment. Right: the DP graph for the same sequence graph and sequence. The dashed grey edges have a cost of 1, and the solid black edges have a cost of 0. The source node 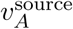 is at the top. The distance from the source node to any node is equal to the score in the DP matrix.

#### Definition 4

(Alignment graph). Given a string *s ∈* Σ^*m*^ and sequence graph *G* = (*V, E, σ*) with *V* = *{v*_1_, …, *v*_*n*_}, we define the corresponding *alignment graph G*_*A*_ through a node set *V*_*A*_:= *V ×* {1, …, |*s*|} and a weight function

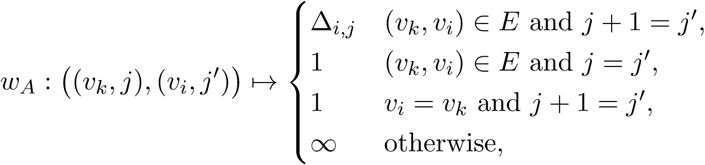

where an edge exists between two nodes from *V*_*A*_ if the corresponding weight is finite. Additionally, we add a special node 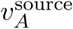 and ∆_*i,*1_-weight edges from 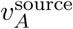 to every node (*v_i_,* 1).

#### Theorem 1

(Existence). For any sequence graph *G* = (*V, E, σ*) and sequence *s ∈* Σ^*m*^, there exists an assignment of cell scores *C*_*i,j*_ that satisfies the recurrence given in Definition 3.

#### Proof.

We consider the corresponding alignment graph. We observe that none of its edges can go “up”, that is, for any edge (*v_k_, j*) *→* (*v_i_, j′*), we always have *j ≤ j′* by Definition 4. Hence, all cycles will be among nodes in the same “row” (i.e. for the same value of *j*). Such “horizontal” edges with *j* = *j′* have a weight of 1 by definition of the weights. Together, this implies that all cycles have a positive, non-zero cost. That, in turn, implies that the minimum distance from the source node 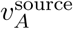 to any other node is well-defined and unique. We set *C*_*i,j*_ to the minimum distance from 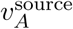 to (*v_i_, j*). This assignment of *C*_*i,j*_ values satisfies the recurrence in Definition 3, which follows immediately by induction since weights in Definition 4 mirror Recurrence (1).

#### Theorem 2

(Uniqueness). For any sequence graph *G* = (*V, E, σ*) and sequence *s ∈* Σ^*m*^, there is exactly one assignment of cell scores *C*_*i,j*_ that satisfies Definition 3.

#### Proof.

The existence of a solution was established in Theorem 1 and we denote the cell scores given by the shortest paths in the corresponding alignment graph by *C*_*i,j*_. Suppose, for the sake of contradiction, that there exists a different assignment 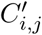 for *i ∈* {*i*, …, |*V* |} and *j ∈* {1, …, |*s*|}. Let *i′* and *j′* be indices such that 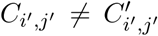. Now consider a shortest path from 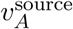 to (*i′ j′*), corresponding to a sequence of nodes 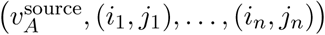 with (*i*_*n*_, *j*_*n*_) = (*i′, j′*). Let *k* be the smallest index such that 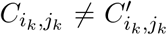. Note that *k >* 1 because the initialization of the first row of the DP table (Definition 3) is identical to the choice of weights of edges from the source node 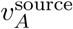 to the first row of vertices (Definition 4), and hence 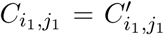. If 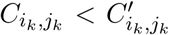, then 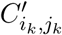 violates Recurrence (1), because Recurrence (1) minimizes over all possible predecessor cells, which correspond exactly to the incident nodes of (*i*_*k*_, *j*_*k*_) in the alignment graph and the term in Recurrence (1) containing 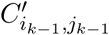 hence gives rise to a smaller value—a contradiction. If 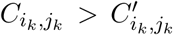, then 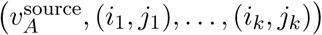 cannot be a shortest path from 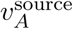 to (*i*_*k*_, *j*_*k*_), because a shorter path can be obtained by following the “backtrace” through the DP table, i.e. by following the sequence of cells given by the minima picked in Recurrence (1).

The proof of Theorem 1 gives us a simple and immediate way to solve Problem 1 by constructing the alignment graph and using Dijkstra’s algorithm [6] to find the shortest path from the source to any node in the bottom row. Using a Fibonacci heap as the priority queue, this requires *O|E|m* + *|V|m* log(*|V|m*) time and *O*(*|V|m*) memory. This strategy comes with the advantage that we can stop the search as soon as it reaches the bottom row. Then the algorithm will have explored only the nodes with a score less than or equal to the optimal alignment score, which can be far smaller than the full matrix for exact or close to exact matches. This method has been used by the hybridSPAdes assembler [1] for mapping long reads to assembly graphs. However, both worst-case runtime and memory requirements of this approach are not optimal, as we show below.

### 2.2 Vertical and Horizontal Properties

Ukkonen proved the *vertical property* and the *horizontal property* in 1985 [27], which state that for the standard semi-global alignment with unit costs, the difference between two vertically or horizontally neighboring cells in the DP matrix is in the range {−1, 0, +1}. This observation is the basis for various optimizations, including Myers’ bit-parallel algorithm [16], which enables a speedup from *O*(*mn*) to 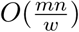 for machines with *w*-bit words. Here we show that the vertical property holds for the graph formalization and that the horizontal property holds for at least one in-neighbor of each cell. These properties are the basis for an algorithm that is asymptotically faster then Dijkstra’s.

#### Theorem 3

(Vertical property). The score difference between any two vertically adjacent cells *C*_*i,j*_ and *C*_*i,j*−1_ is at most one, that is, *C*_*i,j*_ − *C*_*i,j*−1_ *∈* {−1, 0, 1} for all *i ∈* {1, …, |*V*|} and *j ∈* {2, …, |*s*|}.

*Proof.* It is clear from Recurrence 1 that *C*_*i,j*_ − *C*_*i,j*−1_ *≤* 1. Next, we have to prove the bound *C*_*i,j*_ −*C*_*i,j*−1_ ≥ −1 or, equivalently, *C*_*i,j*−1_ ≤ *C*_*i,j*_ +1. We distinguish three cases, based on which of the three terms terms in Recurrence 1 takes a minimum (note that more than one term can have the minimum value).

*Case 1 (vertical, C*_*i,j*_ = *C*_i,j−1_ + 1). This case immediately implies *C*_*i,j*_ − *C*_*i,j*−1_ ≥ −1.

*Case 2 (diagonal, C*_*i,j*_ = *C*_*k,j*−1_ + ∆_*i,j*_ for 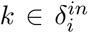). By Recurrence (1), we have *C*_*i,j*−1_ ≤ *C*_*k,j*−1_ + 1. Therefore, *C*_*i,j*−1_ ≤ *C*_*k,j*−1_ + 1 = *C*_*i,j*_ − ∆_*i,j*_ + 1 by the assumption of Case 2. It follows *C*_*i,j*_ ≥ *C*_*i,j*−1_ + ∆_*i,j*_ − 1, which implies *C*_*i,j*_ − *C*_*i,j*−1_ ≥ −1 since ∆_*i,j*_ ≥ 0 by definition.

*Case 3 (horizontal, C*_*i,j*_ = *C*_*k,j*_ + 1 *for* 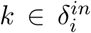). Our proof for this case is by induction on values of the cells in a row. That is, to prove the claim *C*_*i,j*−1_ ≤ *C*_*i,j*_ + 1 for cell *C*_*i,j*_, we assume that it holds for all cells in the same row with smaller value, i.e. for all cells *C*_*i′,j*_ with *C*_*i′,j*_ < *C*_*i,j*_. Note that the claim holds for all cells with minimum value in each row, because they are covered by Case 1 or Case 2, but cannot fall under Case 3. By the induction hypothesis, we have *C*_*k,j*−1_ ≤ *C*_*k,j*_ +1 and, by the assumption of Case 3, we get *C*_*k,j*−1_ ≤ *C*_*i,j*_. By Recurrence 1, we have *C*_*i,j*−1_ ≤ *C*_*k,j*−1_ + 1. Together, this implies that *C*_*i,j*−1_ ≤ *C*_*k,j*−1_ + 1 ≤ *C*_*i,j*_ + 1.

#### Theorem 4

(Horizontal property). For every cell *C*_*i,j*_, whose in-neighbor set is not empty, there is an in-neighbor *C*_*k,j*_ such that the score difference is at most one, that is, there exists a 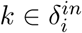 such that *C*_*i,j*_ − *C*_*k,j*_ ∈ {−1, 0, 1}.

#### Proof.

From Recurrence 1 it is clear that the upper bound *C*_*i,j*_ − *C*_*k,j*_ ≤ 1 holds for all in-neighbors. Next we will prove that lower bound 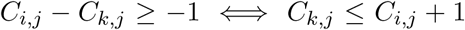 holds for at least one in-neighbor. Like in the proof of Theorem 3, we distinguish three cases.

*Case 1 (horizontal, C*_*i,j*_ = *C*_*k,j*_ + 1 *for* 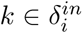). Immediately implies the bound.

*Case 2 (diagonal, C*_*i,j*_ = *C*_*k,j*−1_ + ∆_*i,j*_ for 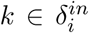). By Theorem 3, the vertical property *C*_*k,j*_ ≤ *C*_*k,j*−1_ + 1 holds. Combining this with *C*_*i,j*_ = *C*_*k,j*−1_ + ∆_*i,j*_ (assumption of Case 2) yields *C*_*k,j*_ ≤ *C*_*i,j*_ − ∆_*i,j*_ + 1. Since ∆_*i,j*_ ≥ 0, this implies the claim.

*Case 3 (vertical, C*_*i,j*_ = *C*_*i,j*−1_ + 1). We prove this case by induction on the row index *j*. For *j* = 1, we have *C*_*i,j*_ ∈ {0, 1} for all *i* by Definition 3 and hence the claim holds. For *j* > 1, the vertical property *C*_*k,j*_ ≤ *C*_*k,j*− 1_ + 1 holds for all *k* by Theorem 3. By the induction hypothesis, there exists a *k* such that *C*_*k,j*−1_ ≤ *C*_*i,j*−1_ + 1, which yields *C*_*k,j*−1_ ≤ *C*_*i,j*_ by plugging in the assumption of Case 3. Together with the vertical property, we get *C*_*k,j*_ ≤ *C*_*i,j*_ + 1.

#### Partial Recurrence Values

The horizontal and vertical properties will be important to build our algorithm, but we need another observation on values of a *partial recurrence*. The idea is to consider the values we obtain when omiting the “horizontal” term “*C*_*k,j*_+1” from Recurrence (1). The next lemma establishes that the values obtained this way are either correct, that is, equal to the values for full recurrence, or too large by exactly 1.

##### Lemma 5

(Partial recurrence values). Let a sequence graph *G* = (*V, E, σ*), a sequence *s ∈* Σ^*m*^, and values *C*_*i,j*_ for *i* ∈ {1, …, |*V* |} and *j ∈* {1, …, *m*} that satisfy Definition 3 be given. The partial recurrence values

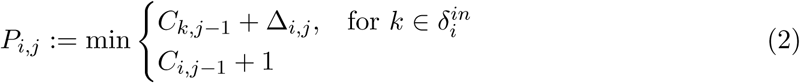

satisfy *P*_*i,j*_ − *C*_*i,j*_ ∈ {0, 1} for all *i ∈* {1, …, |*V* |} and *j ∈* {2, …, *m*}.

##### Proof.

Recurrence (1) and the definition of *P*_*i,j*_ immediately imply that *P*_*i,j*_ −*C*_*i,j*_ ≥ 0. To prove *P*_*i,j*_ − *C*_*i,j*_ ≤ 1, suppose there existed *i* and *j* such that *P*_*i,j*_ − *C*_*i,j*_ > 1. By their definitions, *P*_*i,j*_ and *C*_*i,j*_ can only differ when the minimum in Recurrence (1) came from the “horizontal” term *C*_*k,j*_ + 1. Therefore, there exists 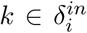 such that *C*_*k,j*_ + 1 = *C*_*i,j*_. By the vertical property (Theorem 3), we get *C*_*k,j*−1_ ≤ *C*_*k,j*_ + 1, implying that *C*_*k,j*−1_ ≤ *C*_*i,j*_. Since ∆_*i,j*_ ≤ 1, this means that *P*_*i,j*_ ≤ *C*_*k,j−1*_ + 1 = *C*_*i,j*_ + 1, a contradiction.

### 2.3 Algorithm for Semi-Global Sequence-to-Graph Alignment with Unit Costs

We now devise an algorithm to compute the optimal alignment cost in *O*(*|E|m*) time and *O*(*|V |*) space. To this end, we process the matrix row by row. To compute the scores in a row, we first apply the “partial recurrence” as given in Lemma 5—that is, we only use the “diagonal” and “vertical” terms of Recurrence 1—resulting in “almost” correct scores. We then process the cells in that row ordered by their value and apply the “horizontal term”. We prove this process to lead to the correct values irrespective of the cyclic dependencies.

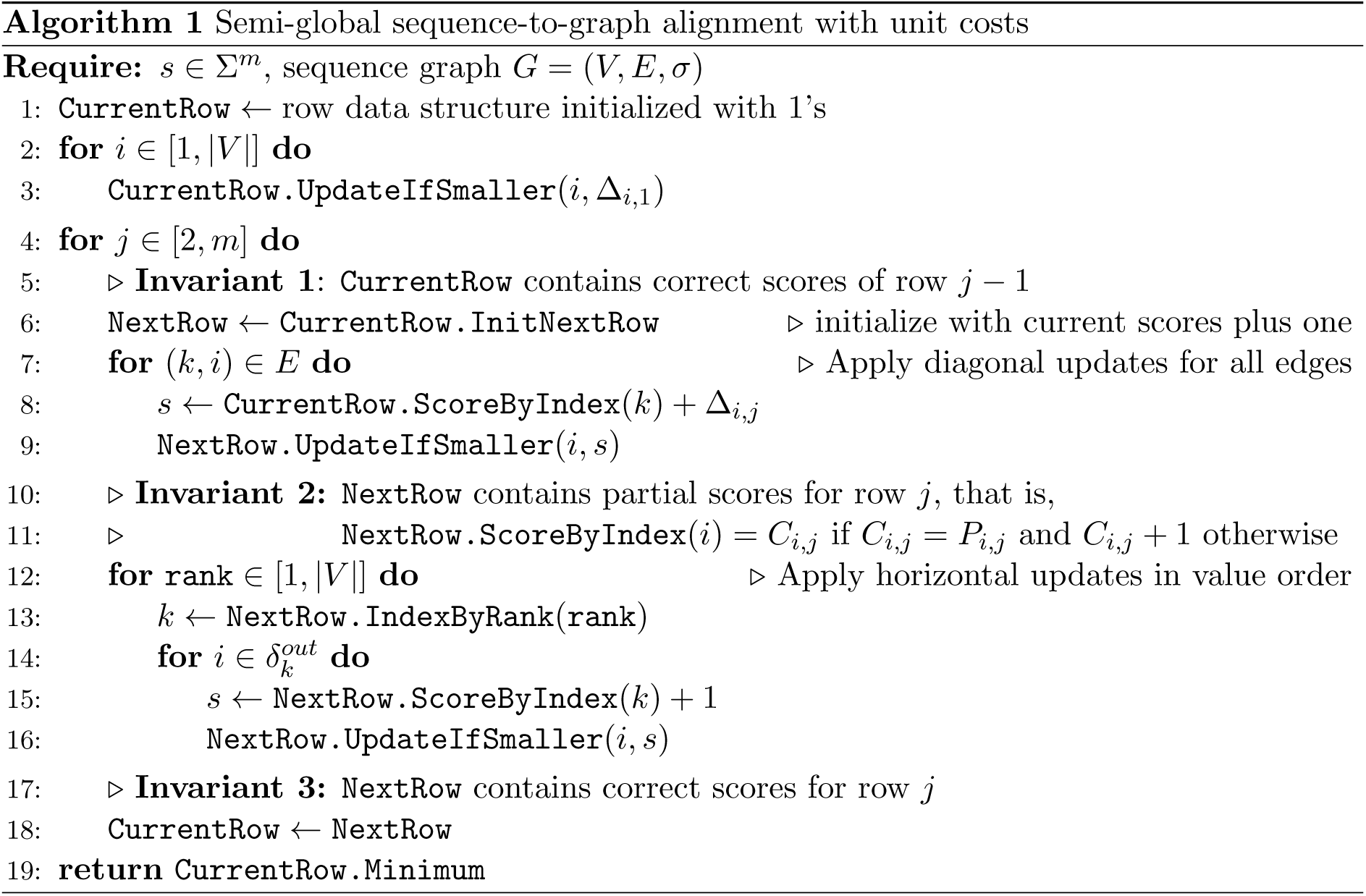

Pseudocode for our approach is given in Algorithm 1. It makes use of an auxiliary data structure that represents one row of the DP matrix and supports the following operations:

- `ScoreByIndex`(*i*): Return the value of column *i* in the represented row.
- `UpdateIfSmaller`(*i, s*): If *s* is smaller then the current the value of column *i*, then set the value to *s*, otherwise do nothing.
- `IndexByRank(rank)`: Return the index *i* of the column at the given rank with respect to an ordering by value. Ties are resolved arbitrarily, but in a way such that subsequent calls to `IndexByRank` give consistent results (i.e. calling it for all ranks results in all indices).
- `InitNextRow`: Create a new data structure instance to represent the next row and initialize all values to the current values plus one.

Before providing details on how to efficiently implement such a data structure, we verify that Algorithm 1 is correct.

#### Theorem 6

(Correctness of Algorithm 1). For a sequence graph *G* = (*V, E, σ*) and a sequence *s ∈* Σ^*m*^, Algorithm 1 computes the minimum edit distance min_*p*_ *d*(*p, s*) over all paths *p* in *G*.

#### Proof.

Recurrence 1 directly corresponds to the possible edit operations. Therefore, it is sufficient to show that Algorithm 1 correctly computes values *C_i_,|m|* that satisfy Definition 3. To this end, let us look at the three invariants in the pseudocode. Lines 2 to 3 initialize the values for the first row exactly as prescribed in Definition 3. Therefore, Invariant 1 is true for the first row. For later rows, Invariant 1 holds for row *n* if Invariant 3 holds for the row *n −* 1.

Now, consider Invariant 2. Line 6 initializes the values as *C*_*i,j*_ = *C*_*i,j*−1_ + 1. This is equivalent to initializing *C*_*i,j*_ via the “vertical” recurrence term. The loop in lines 7 to 9 updates each cell via the “diagonal” recurrence terms. We note that after running lines 6 to 9, each “vertical” and “diagonal” term has been checked. This is equivalent to calculating the partial recurrence score, so Invariant 2 for row *n* is true if Invariant 1 is true for row *n*.

For Invariant 3, we have to prove that the loop in lines 12 to 16 updates the partial recurrence score to the true recurrence score. Line 13 retrieves the index *k* with a given rank. At that point, the entry in NextRow at index *k* is guaranteed to be correct, i.e. to be equal to *C_k_, j*, which we show by contradiction. Assume it was equal to *C*_*k,j*_ + 1, which is the only other option by Lemma 5. Then, there must exist a node index *k′*, such that *C*_*k,j*_ = *C*_*k′,j*_ + 1; i.e. a DP cell in the same row giving rise to a horizontal update. Again by Lemma 5, the current row entry at index *k′* can either be *C*_*k′,j*_ or *C*_*k′,j*_ + 1, both of which are strictly smaller than the current entry at index *k*, which is equal to *C*_*k,j*_ + 1. Therefore, index *k′* must have been processed already and updated its out-neighbors in Line 16. Which contradicts a value of *C*_*k,j*_ + 1 at index *k*.

So in Line 13, the value at index *k* is correct. Furthermore, Line 13 is guaranteed to process all indices because even though the ranks can be updated in Line 16, none can be skipped because the updated values computed in Line 15 are larger than the value at index *k* (and thus retain a larger rank after an update). Thus, the scores will be correct and Invariant 3 is true for row *n* if Invariants 1 and 2 are true for row *n*. Since Invariant 1 is true for the first row, all invariants are true by induction.

#### Row Data Structure

To achieve fast runtimes of Algorithm 1, we need an efficient data structure to accommodate the cost values in a row and their order. Below, we describe a data structure that can be initialized in *O*(*|V |*) time and supports the `ScoreByIndex` and `IndexByRank` operations in *O*(1) time, `InitNextRow` in *O*(*|V |*) time, and `UpdateIfSmaller` in *O*(*d*) time, where *d* is the difference between the target value and the current value. Internally, we maintain five arrays:

- `value_space`: the universe of possible values in that row, sorted by value,
- `order`: column indices sorted ascending by value,
- `rank`: inverse of the order array, that is, a column’s position in the sorted array. Therefore, `order` [`rank`[*i*]] = *i* and `rank` [`order`[*i*]] = *i*,
- `vi_by_index`: the value index (vi) with respect to `value_space` of the column at a given index *i*, i.e. `value_space`[`vi_by_index`[*i*]] is value of column *i*,
- `min_index`: for each entry in `value_space`, the array `min_index` gives the minimum rank such that the value at this rank is larger than the corresponding value, that is, `vi_by_index`[`order`[`min_index`[*vi*] *−* 1] ≤ *vi* and `vi_by_index[order[min_index`[*vi*]] > *vi*.

The arrays `order`, `rank`, and `vi_by_index` have a size of *|V |*, while `value_space` and `min_index` can have a different size, i.e. |`value_space`| = |`min_index`| ≠ *|V |*. Upon initialization for the first row `value_space` = [0, 1] because 0 and 1 are the only possible values in the first row. When initializating a subsequent row via the `InitNextRow` operation, we employ the vertical property to deduce the universe of possible values: only values present in the current row, plus/minus one, can be present in the next row. Therefore, `value_space` and `min_index` have a size of at most 3*|V |* and the whole data structure needs *O*(*|V |*) space. Given the semantics of theses five arrays, implementing all operations in the runtimes specified above is straighforward and we defer the details to Appendix A.1.

#### Runtime

First, we note that the next row is initialized in Line 6 to scores one higher than the current row. By the vertical property in Theorem 3, the difference between the initialized score and the lowest possible score is two. This means that even though the `UpdateIfSmaller` operation is linear to the score difference, the score difference is at most 2 so all calls to `UpdateIfSmaller` run in constant time. Initialization in line 6 takes *O*(*V*) time. The loop in lines 7 to 9 runs once for each edge. The loop in lines 12 to 16 runs once for each node, and the inner loop iterates over each neighbor, meaning that the innermost block runs once for each edge. So the runtime for each row is *O*(*V* + *E*).

However, we can apply a transformation on the graph which removes enough nodes to guarantee that *|V | ∈ O*(*E*) without affecting the correctness. Since the score for each node *i* with zero in-degree will be *C*_*i,j*_ = *j −* 1 + ∆_*i,*1_, we can merge all nodes with an in-degree 0 and the same node label. The only remaining nodes will be nodes with an incoming edge, plus at most one node for each character in the alphabet, so the number of resulting nodes will be *|V′| ≤ |E|* + |Σ|. Assuming a constant alphabet, this means that *|V ′| ∈ O*(*E*), therefore the runtime per row is *O*(*E*) regardless of the original number of nodes. This transformation means that we have to iterate over the entire graph, so the runtime of the transformation is *O*(*V* + *E*). Since there are *m* rows, the final runtime is *O*(*V* + *mE*).

#### Space Consumption and Backtrace

In case only the minimum edit distance is to be computed, we only need to store the current row and the next row at any point in time and the algorithm uses *O*(*|V |*) space. If the alignment is desired, we can produce a backtrace by storing the origin of each minimum exactly as done for classic sequence-to-sequence alignment. We use the “checkpoint-and-recalculate” strategy [8] and only store every 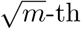 row. This leads to a space consumption of 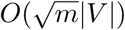 and no overhead in (asymptotic) runtime.

## 3 Generalizing Alignment Costs

### Arbitrary match costs

Existence and uniqueness (theorems 1 and 2) do not depend specifically on Recurrence 1 having unit costs. The only requirement on costs is that the “horizontal” cost is positive and non-zero. However, the other costs, match, mismatch, and insertion, cannot form cycles in the alignment graph, and so can be any real numbers, including zero and negative numbers. This means that we can also use different, non-unit costs such as substitution matrices [20]. It also means that the problem can be rephrased as a match score maximization problem instead of a cost minimization. For score maximization, we have to multiply the scores by *−*1 and then find the minimum score; this is trivially the same as finding the maximum score.

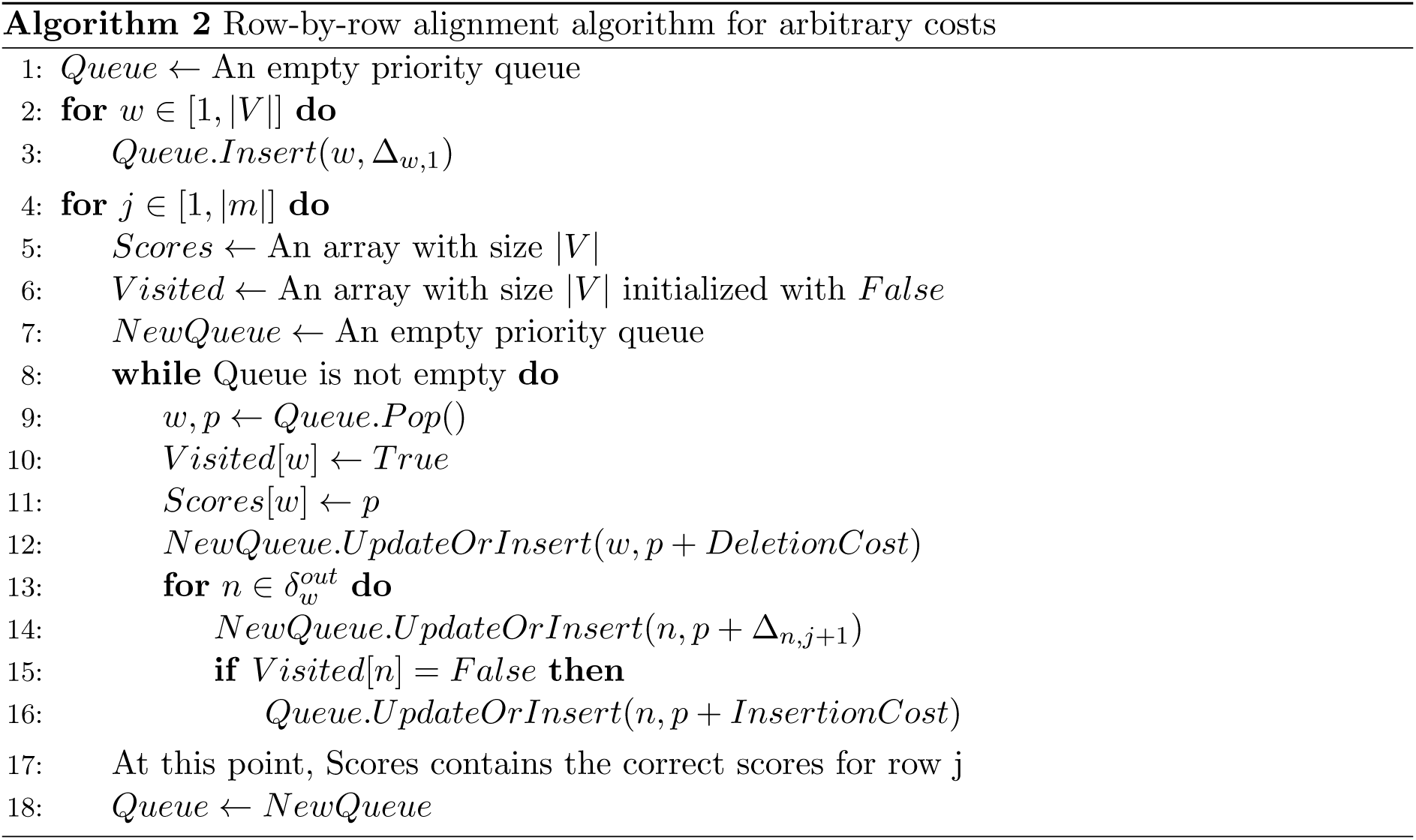

Algorithm 2 shows the pseudocode for the arbitrary cost algorithm. We use a priority queue to make sure that the cells are processed in ascending order by score. We assume that our priority queue has an *UpdateOrInsert* operation, which updates an existing item’s priority if the given priority is smaller than the current priority, or inserts a nonexistant item with the given priority. We use a node’s score as its priority, so the nodes are processed in ascending score order. When processing a node, the priorities of its out-neighbors in the alignment graph are updated. This might update a node in the same row as the current node. However, since the only edge into the same row is the deletion cost, and since deletion cost is positive and non-zero, the out-neighbor has a higher score both before and after the update. For each row, each node is inserted into and popped from the queue exactly once and updated up to *E* times. After a node has been processed, the if condition in Line 15 stops it from being inserted into the queue again. Therefore each node is in the queue exactly once, and the while loop runs *V* times. The for loop in Line 13 processes each edge once. Each edge causes a priority lookup and possibly a priority decrement. The runtime of the algorithm depends on the priority queue implementation used. For each row, there are *O*(*E*) score lookups, score updates and insertions; and *O*(*V*) pops. Using a Fibonacci heap, the score lookup, update and insertion are *O*(1) and pop is *O*(log *n*). This leads to a runtime *O*(*|E|m* + *|V |m* log *|V |*) with *O*(*|V |*) memory usage.

### Affine gap penalty

In the *affine gap cost* model, starting a gap and extending a gap have different costs. Formally, we define the recurrence:

#### Definition 5

(Recurrence for affine gap cost SG^2^A). Set *C*_*i*,1_ = ∆_*i,*1_ for all *i ∈* {1, …, |*V*|}. And define

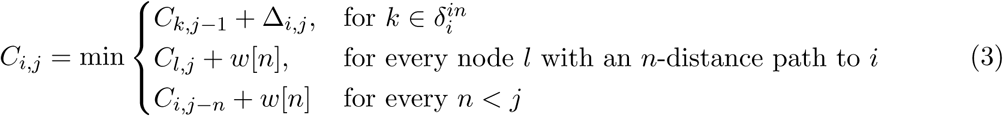

where ∆_*i,j*_ the mismatch penalty between node character *σ*(*v*_*i*_) and sequence character *s*_*j*_, which is an arbitrary real number, and *w*[*n*] is the cost of an affine gap with length *n*; *w*[*n*] = *a*+(*n−*1)*b* for some constants *a* > 0 and *b* > 0.

For the standard DP formulation, this is typically implemented with separate matrices for insertions and deletions, and adding terms to the recurrence for transitioning between the normal DP matrix and the affine gap matrices [7]. The same idea can be used for the alignment graph approach. The alignment graph is now split into three subgraphs: *match subgraph*, *insertion subgraph* and *deletion subgraph*. Now, instead of having vertical and horizontal edges in the match subgraph for insertions and deletions, these edges connect from the node in the match subgraph to the corresponding neighbor node in the insertion and deletion subgraphs. The cost of the edges from the match subgraph to the insertion or deletion subgraph is the gap start cost, and the cost of edges within the insertion and deletion subgraphs is the gap extension cost. Finally, we add an edge from every node in the insertion and deletion subgraphs to the corresponding neighbor node in the match subgraph. Formally, the alignment graph for affine gap costs is defined as:

#### Definition 6

(Affine gap alignment graph). Given constants *a* > 0, *b* > 0, a string *s ∈* Σ^*m*^ and sequence graph *G* = (*V, E, σ*) with *V* = *{v*_1_, …, *v*_*n*_}, we define the corresponding *affine gap alignment graph G*_*A*_ through the node set *V*_*A*_:= *V* × {1, …, |*s*|} × {*M, I, D*}, consisting of the *match subgraph* (M), *insertion subgraph* (I) and *deletion subgraph* (D), and a weight function

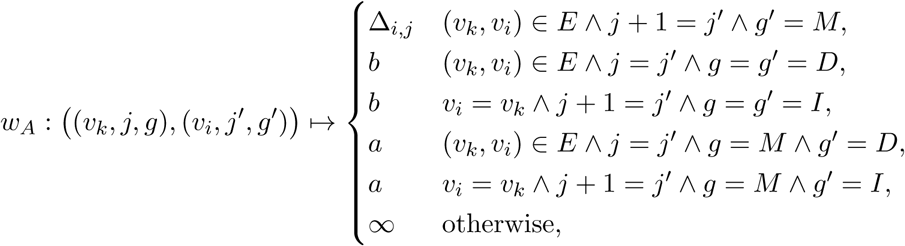

where an edge exists between two nodes from *V*_*A*_ if the corresponding weight is finite. Additionally, we add a special node 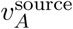 and add ∆_*i,*1_-weight edges from 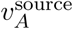 to every node (*v_i_,* 1, *M*) for *i ∈* {1, …, *n*}.

Figure 2 shows an example of an affine gap alignment graph. Now, an insertion (or deletion) is represented as a path that starts in the match subgraph, moves to the insertion (or deletion) subgraph, and then back to the match subgraph at a different node. The unit cost algorithm cannot be used for affine gaps since the gap start cost is typically higher than one. We note that there are no negative or zero-cost cycles, and every edge is either between nodes in the same row or directly adjacent rows, so the arbitrary cost algorithm can be used for the affine gap alignment graph with only slight modifications and with the same runtime and space requirements.

**Figure 2:**
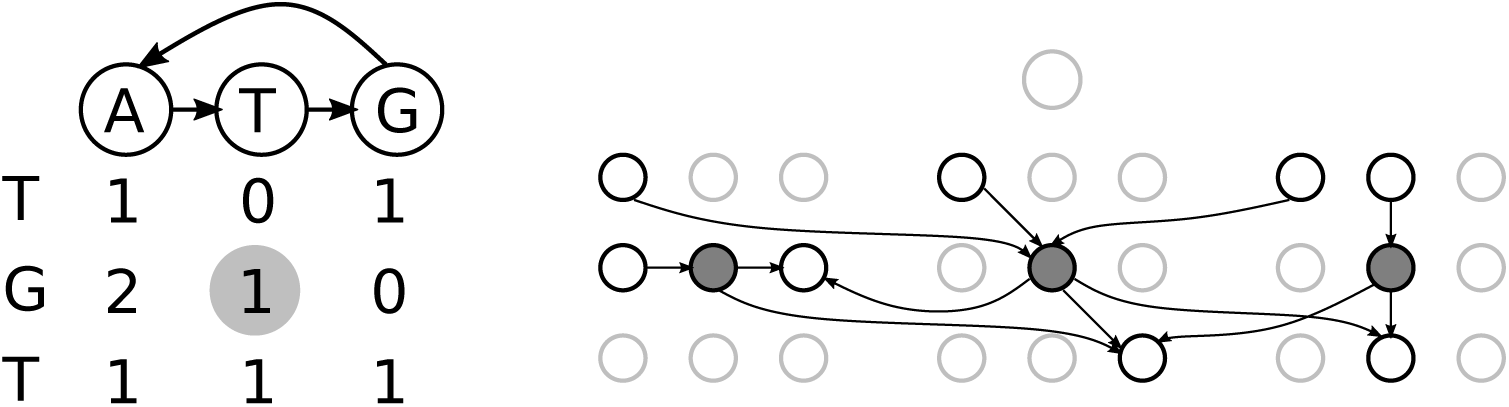
Left: the DP matrix between a sequence graph (top) and a sequence (left). Right: the affine gap alignment graph of the sequence graph and sequence. The three grey nodes represent the middle cell in the DP matrix. For clarity, only edges connecting to or from the grey nodes are shown. The graph is divided into three subgraphs, the match subgraph (middle), deletion subgraph (left) and insertion subgraph (right).

## 4 Discussion

In this paper, we establish important properties of alignments of sequences to directed node-labeled graphs. We prove existence and uniqueness of a solution, horizontal and vertical properties, as well as properties of partial recurrence values. These results enable an *O*(*V* + *mE*) algorithm for aligning a sequence to an arbitrary directed graph with unit costs. To our knowledge, this is the fastest published sequence-to-graph alignment algorithm that can be applied to cyclic graphs. Its runtime is optimal in the sense that *O*(*n*^2*−ϵ*^) algorithms for the special case of sequence-to-sequence alignment are unlikely to exist. It is open, however, whether faster algorithms for special graph classes are possible, for instance for graphs with more edges than nodes, i.e. with *E* in *ω*(*V*). We have also presented an *O*(*mE* + *V_m_* log *V*) algorithm for optimally aligning a sequence to a graph with affine gap costs and arbitrary match costs. We emphasize that the runtimes of both algorithm, for unit costs and affine gap costs, depend solely on the size of the graph and the length of the sequence, but not on shape of the graph.

Many other flavors of sequence-labeled graphs exist and are used in bioinformatics [25, 19], such as *de Bruijn graphs*, *overlap graphs* and *variation graphs* [19]. These graphs are often considered in their bi-directed version [15, 19], so that each node likewise represents the reverse complement sequence. Although we lack the space to expand on this, we note that all these graphs can likewise be handled in our framework by suitably transforming these graphs into node-labeled directed graphs, without an asymptotic increase in the graph size (except for splitting possible multi-character node/edge labels into multiple nodes labeled by single characters).

The algorithms we present here are not directly suited for aligning long reads to genome-scale graphs. Nonetheless, the theory we provide in this paper provides an excellent basis for engineering algorihms that are fast in practice. To this end, we are currently working on adapting the classic ideas of seeded alignments [3], banded alignment [26], and bit-parallelism [17] to sequence-to-graph alignment.

## Acknowledgments

We thank Shilpa Garg for inspiring discussions and for feedback on a draft version.

## A Appendix

### A.1 Implementation Details of Row Data Structure

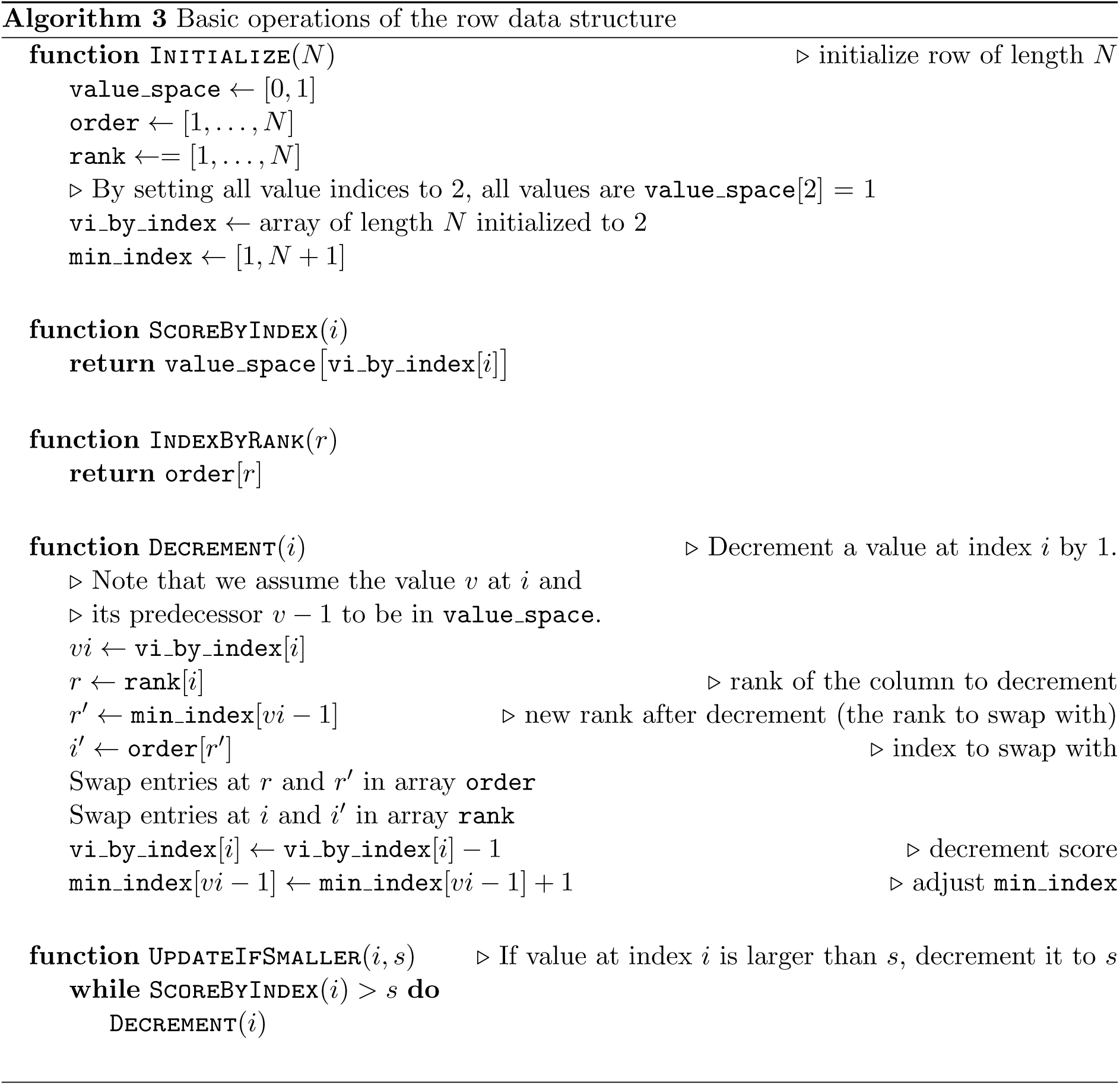

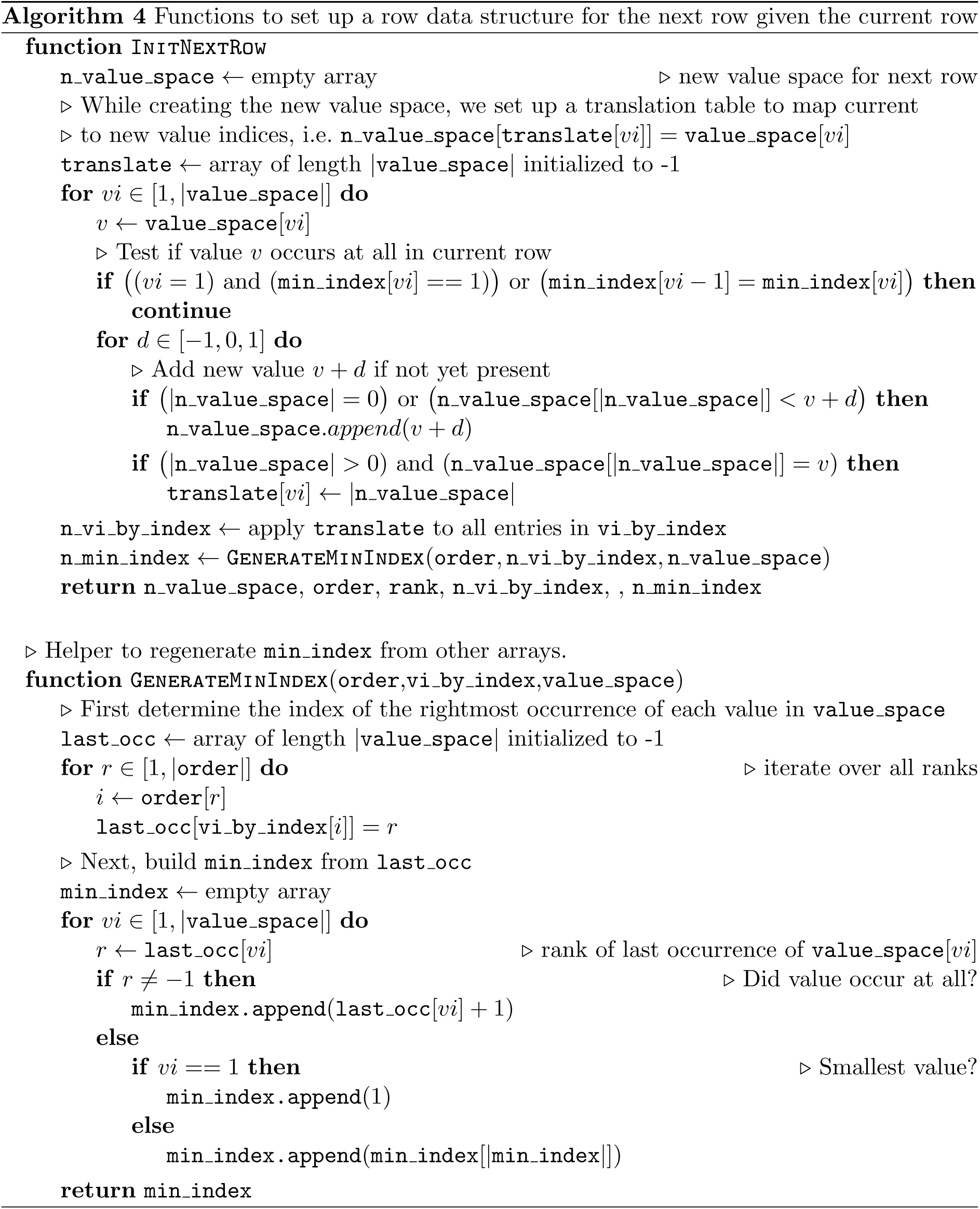

## References

[1] Antipov, D., Korobeynikov, A., McLean, J.S., Pevzner, P.A.: hybridSPAdes: an algorithm for hybrid assembly of short and long reads. Bioinformatics 32(7), 1009–1015 (2016), +http://dx.doi.org/10.1093/bioinformatics/btv688

[2] Backurs, A., Indyk, P.: Edit distance cannot be computed in strongly subquadratic time (unless SETH is false). In: Proceedings of the Forty-seventh Annual ACM Symposium on Theory of Computing. pp. 51–58. STOC ’15, ACM, New York, NY, USA (2015)

[3] Burkhardt, S., Crauser, A., Ferragina, P., Lenhof, H.P., Rivals, E., Vingron, M.: q -gram based database searching using a suffix array (QUASAR). In: Proceedings of the third annual international conference on Computational molecular biology. pp. 77–83. ACM (Apr 1999)

[4] Canzar, S., Slazberg, S.L.: Short Read Mapping: An Algorithmic Tour. Proc. IEEE PP(99), 1–23 (2015)

[5] Compeau, P.E.C., Pevzner, P.A., Tesler, G.: How to apply de bruijn graphs to genome assembly. Nat. Biotechnol. 29(11), 987–991 (8 Nov 2011)

[6] Dijkstra, E.W.: A note on two problems in connexion with graphs. Numerische Mathematik 1(1), 269–271 (Dec 1959), https://doi.org/10.1007/BF01386390

[7] Gotoh, O.: An improved algorithm for matching biological sequences. Journal of molecular biology 162(3), 705–708 (1982)

[8] Grice, J.A., Hughey, R., Speck, D.: Reduced space sequence alignment. Bioinformatics 13(1), 45–53 (1997)

[9] Huang, L., Popic, V., Batzoglou, S.: Short read alignment with populations of genomes. Bioinformatics 29(13), i361–i370 (Jul 2013), http://bioinformatics.oxfordjournals.org/content/29/13/i361

[10] Kehr, B., Trappe, K., Holtgrewe, M., Reinert, K.: Genome alignment with graph data structures: a comparison. BMC Bioinformatics 15(1), 99 (Apr 2014), http://www.biomedcentral.com/1471-2105/15/99/abstract

[11] Lee, C., Grasso, C., Sharlow, M.F.: Multiple sequence alignment using partial order graphs. Bioinformatics 18(3), 452–464 (2002), +http://dx.doi.org/10.1093/bioinformatics/18.3.452

[12] Limasset, A., Cazaux, B., Rivals, E., Peterlongo, P.: Read mapping on de bruijn graphs. BMC Bioinformatics 17(1), 237 (16 Jun 2016)

[13] Miller, J.R., Koren, S., Sutton, G.: Assembly algorithms for next-generation sequencing data. Genomics 95(6), 315–327 (Jun 2010)

[14] Myers, E.W.: An overview of sequence comparison algorithms in molecular biology. Tech. Rep. 91-29, Department of Computer Science, University of Arizona (1991)

[15] Myers, E.W.: Toward simplifying and accurately formulating fragment assembly. J. Comput. Biol. 2(2), 275–290 (1995)

[16] Myers, G.: A fast bit-vector algorithm for approximate string matching based on dynamic programming. J. ACM 46(3), 395–415 (May 1999), http://doi.acm.org/10.1145/316542.316550

[17] Myers, G.: A fast bit-vector algorithm for approximate string matching based on dynamic programming. J. ACM 46(3), 395–415 (May 1999)

[18] Needleman, S.B., Wunsch, C.D.: A general method applicable to the search for similarities in the amino acid sequence of two proteins. Journal of Molecular Biology 48(3), 443–453 (1970), http://www.sciencedirect.com/science/article/pii/0022283670900574

[19] Paten, B., Novak, A.M., Eizenga, J.M., Garrison, E.: Genome graphs and the evolution of genome inference. Genome research 27(5), 665–676 (2017)

[20] Pearson, W.R.: Selecting the right similarity-scoring matrix. Current protocols in bioinformatics pp. 3–5 (2013)

[21] Salmela, L., Rivals, E.: Lordec: accurate and efficient long read error correction. Bioinformatics 30(24), 3506–3514 (2014), +http://dx.doi.org/10.1093/bioinformatics/btu538

[22] Schneeberger, K., Hagmann, J., Ossowski, S., Warthmann, N., Gesing, S., Kohlbacher, O., Weigel, D.: Simultaneous alignment of short reads against multiple genomes. Genome Biology 10(9), R98 (Sep 2009), http://genomebiology.com/2009/10/9/R98/abstract

[23] Sellers, P.H.: The theory and computation of evolutionary distances: Pattern recognition. J. Algorithm. Comput. Technol. 1(4), 359–373 (Dec 1980)

[24] Smith, T.F., Waterman, M.S.: Identification of common molecular subsequences. Journal of molecular biology 147(1), 195–197 (1981)

[25] The Computational Pan-Genomics Consortium: Computational pan-genomics: status, promises and challenges. Briefings in Bioinformatics p. bbw089 (Oct 2016), http://bib.oxfordjournals.org/content/early/2016/10/19/bib.bbw089

[26] Ukkonen, E.: Algorithms for approximate string matching. Information and control 64(1–3), 100–118 (1985)

[27] Ukkonen, E.: Finding approximate patterns in strings. Journal of Algorithms 6(1), 132–137 (1985), http://www.sciencedirect.com/science/article/pii/0196677485900239

[28] Vaddadi, K., Sivadasan, N., Tayal, K., Srinivasan, R.: Sequence alignment on directed graphs. bioRxiv (2017), http://www.biorxiv.org/content/early/2017/04/06/124941

